# Cell-type specific arousal-dependent modulation of thalamic activity in the lateral geniculate nucleus

**DOI:** 10.1101/2021.02.17.431649

**Authors:** Benedek Molnár, Péter Sere, Sándor Bordé, Krisztián Koós, Péter Horváth, Magor L. Lőrincz

## Abstract

State dependent thalamocortical activity is important for sensory coding, oscillations and cognition. The lateral geniculate nucleus (LGN) relays visual information to the cortex, but the state dependent spontaneous and visually evoked activity of LGN neurons in awake behaving animals remains controversial. Using a combination of pupillometry, extracellular and intracellular recordings from identified LGN neurons we show that thalamocortical neurons and interneurons are distinctly correlated to arousal forming two complementary coalitions. Intracellular recordings indicated that the membrane potential of LGN TC neurons was tightly correlated to fluctuations in pupil size. Inactivating the corticothalamic feedback to the LGN suppressed the arousal dependency of LGN neurons. Taken together our results show that LGN neuronal membrane potential and action potential output are dynamically linked to arousal dependent brain states in awake mice and this might have important functional implications.

---

Brain states can fluctuate on various timescales and can profoundly influence neural and behavioral responses (McGinley MJ, M Vinck, et al. 2015). This is most apparent when spontaneous neuronal activity and responses to sensory stimuli are compared between states of sleep and wakefulness (Livingstone MS and DH Hubel 1981; Massimini M et al. 2005), but recently, the prominent influence of spontaneous variations within the waking state on both cortical neuronal responses and perceptual abilities has been documented in both humans (Fox MD and ME Raichle 2007) and rodents (Reimer J et al. 2014; McGinley MJ, SV David, et al. 2015; Vinck M et al. 2015). These state dependent activities are reliant upon finely tuned interactions between neocortical and thalamic excitatory and inhibitory neuronal assemblies, strongly influenced by neuromodulatory inputs and lead to various forms of rhythmic activities in thalamo-cortico-thalamic networks (Lorincz ML and AR Adamantidis 2017). Driven by intrinsic and network mechanisms, state dependent thalamic oscillations are thought to possess both a rhythm-regulation function, and a plasticity function important for the faithful sensory information processing during attentive wakefulness (Crunelli V et al. 2018).

Despite the large body of work aiming to identify the cellular and network mechanisms of thalamic low vigilance state oscillations, relatively little is known about the activity of thalamic neurons during brain state changes within the waking state. During relaxed wakefulness LGN TC and interneurons in cats fire phase locked to and contribute to the generation of EEG α (8–13 Hz) rhythms characteristic of inattentive states (Hughes SW et al. 2004; Lorincz ML et al. 2009). Both baseline activity and visual responses are markedly altered between attentive and inattentive states in LGN neurons of immobile rabbits (Bezdudnaya T et al. 2006; Cano M et al. 2006). Interrestingly, mouse LGN neuronal activity was claimed not to be state dependent, as locomotion, a state associated with high arousal (Vinck M *et al*. 2015) and an increase in the gain of visual cortical responses (Niell CM and MP Stryker 2010; Bennett C et al. 2013; Polack PO et al. 2013; Reimer J *et al*. 2014) failed to alter the spontaneous and visually evoked firing rate of LGN neurons (Niell CM and MP Stryker 2010). Despite the lack of α rhythms in rodents, some studies have revealed a pronounced 3-5 Hz oscillation in the mouse visual system, a potential transient analogue of α rhythms (Bennett C et al. 2013; Einstein MC et al. 2017; Senzai Y et al. 2019) suggesting a strong state dependency of the visual thalamocortical system.

To resolve this controversy, we sought to test the arousal dependent activity of single identified LGN neurons and reveal the origin and mechanisms involved. Using a combination of extra- and intracellular recordings from identified LGN neurons in awake behaving mice, pupillometry and pharmacological inactivation we show that: i) spontaneous activity in thalamic neurons is correlated with state transitions in a cell-type specific manner, with TC neurons being positively and LGN interneurons negatively correlated to arousal ii) the membrane potential of LGN neurons correlates with spontaneous fluctuations in pupil diameter providing a mechanistic explanation to the observed phenomenon, consistent with previous studies conducted in the neocortex (Reimer J *et al*. 2014; McGinley MJ, M Vinck, *et al*. 2015; Vinck M *et al*. 2015) and iii) inactivating the corticothalamic feedback from V1 suppresses the state dependence of LGN activity, revealing its origin. These results can have important implications for the state dependent function of thalamo-cortico-thalamic networks.

## MATERIALS AND METHODS

All experimental procedures were performed according to the European Communities Council Directives of 1986 (86/609/EEC) and 2003 (2003/65/CE) for animal research and were approved by the Ethics Committee of the University of Szeged. 36 C57BL/6 mice of either sex aged 2 to 6 months were used in this study.

### Surgical preparation

Mice were anaesthetized with ketamine/xylazine (100 mg/kg and 10 mg/kg, respectively) and mounted in a stereotaxic frame (Model 902, David Kopf Instruments, USA), the skull of the animal exposed and a stainless steel head post cemented over the frontal suture with dental acrylic resin (Pattern Resin LS, GC America, USA). Craniotomy positions were marked at the following stereotaxic coordinates based on The Mouse Brain in Stereotaxic Coordinates (Paxinos & Franklin, 2004): LGN: 2.3 mm posterior; 2.1 mm lateral to Bregma, V_1_: 3.3 mm posterior; 2.5 mm lateral to Bregma.

For postoperative care, mice received Rimadyl (5 mg/kg, intraperitoneally, Pfizer, USA) and an intramuscular injection of Gentamycin (0.1 mg/kg). Following an initial recovery from surgery of at least 5 days mice were handled gently each day for 7 days to reduce excessive stress or anxiety during the recording sessions and head fixation procedure gradually increased. On the day of the recording, small craniotomies (0.8-1 mm) were performed under isoflurane anesthesia (Forane, Abbvie, USA; dose: 1 L/min 1-1.5% isoflurane and 99-98.5% O_2_) using a microdrill at the positions previously marked leaving the dura mater intact. Mineral oil applied on the dural surface prevented dehydration. Mice were then transferred to an *in vivo* electrophysiological recording setup where their head posts were clamped in a custom apparatus and recording sessions started at least 60 minutes following awakening.

### Pupillometry

Pupillometry was conducted with a Genie (Teledyne Dalsa, Canada) high frame rate (300 fps) infrared camera focused on the eye of the animal illuminated with an infrared LED (900 nm, Marubeni Corp).

### In vivo electrophysiology and juxtacellular labeling

Independently mounted single unit and local field potential (LFP) recordings were performed from the LGN and visual cortex, respectively using either glass micropipettes filled with 0.5 M NaCl solution containing with 1.5% w/v Biocytin (Sigma Aldrich, USA, impedance 3-25 MΩ) or Silicon probes (single shank, 32-channel, Neuronexus). A motorized micromanipulator (Burleigh 8200 Inchworm, USA) and an oil hydraulic micromanipulator (MO-10, Narishige) were used for advancing the microelectrodes to the LGN and V1, respectively. The biological signals were pre-amplified with Axon HS-9A headstages (Molecular Devices, USA) and amplified with an Axoclamp 900A amplifier (Molecular Devices) (gain: 100 and 50 for the LGN and V1, respectively) and filtered (0.1 Hz −200 Hz for LFP, 0.3-6 kHz for units). The amplified signals were then digitized with a CED Power3 1401 AD converter (Cambridge Electronic Design) at 30 kHz sampling rate using Spike2 software (Cambridge Electronic Design). Signals recorded with silicon probes were amplified, filtered and digitized using an Intan circuit board (RHD2000, Intan Technologies).

To confirm the morphology and location of the recorded cells as LGN neurons, we performed juxtacellular labeling of LGN cells (n=7) by applying pulses of anodal current (1-3 nA, 500 ms, 50% duty cycle) for 2-5 minutes (Pinault D 1996). At the end of an electrophysiological recording session, overanaesthetized mice were transcardially perfused with cold phosphate-buffered solution (PB, 100 mM, pH: 7.4), followed by 4% paraformaldehyde (PFA), the brain removed and transferred to 4% PFA solution in 100 mM PB and stored at 4°C overnight. Coronal brain sections containing the LGN was sliced to 50 µm slices with a VT1000S vibratome (Leica, Germany). During the histological processing, slices were cryoprotected with 10% and 20% sucrose solution, cells were opened with a freeze-thaw method and TBS-Tween20 to be able to conjugate biocytin in labelled cells with Cy3-Streptavidin (Jackson ImmunoResearch, USA) as secondary antibody. After 2 hours of incubation with Cy3-Streptavidin, the slices were mounted on glass microscope slides and covered with cover slips. A BX60 fluorescent microscope (Olympus, Japan) and a Surveyor software (Objective Imaging, UK) was used to visualize labeled neurons.

### Cortical inactivation with muscimol

For inactivation of the visual cortex we microinjected muscimol, a GABA_A_ receptor agonist (200 nl, 1 mM) through a glass clamp pipette (15 µm pip diameter). The correlation of the LGN neuron firing and pupil diameter was quantified and compared before and after visual cortical inactivation (n=7 neurons).

### Data analysis

Periods of NREM and REM sleep identified by cortical LFP analysis and rhythmic eye movements were excluded from data analysis. Data analysis was performed off-line with custom-written MATLAB and ImageJ routines. Pupillometry was carried out off-line on recoded AVI files with a custom ImageJ plug-in, which can generally be used for detection of circular objects through minimizing an energy function by a variational framework. LGN neurons were identified as TC or interneurons based on either their morphology (n=7 TC neurons labeled), action potential duration and by the presence, or lack of LTCP mediated burst firing, respectively during periods of quiet wakefulness or NREM sleep.

## RESULTS

We performed extracellular and intracellular recordings of identified thalamic neurons with simultaneous local field potential in the primary visual cortex (V_1_ LFP) and pupilometry in awake head-restrained mice to reveal the brain state dependent activity of thalamic neurons in the LGN.

### The baseline activity of most LGN neurons correlates with brain states

States of quiet wakefulness (QW) were characterized by large-amplitude slow V1 LFP fluctuations, periods of active wakefulness (AW) by small-amplitude fast fluctuations in the LFP recorded in V1 (Fig. 1). The low-frequency spectral power (0.5–5 Hz) during QW was significantly larger than during AW, but high-frequency spectral power (30–80 Hz) was larger during AW than QW (p < 0.001 for both cases, ANOVA for repeated measures). State transitions defined by the amplitude of the LFP recorded from V_1_ were accompanied by prominent changes in the activity of this TC neuron, such as QW to AW transitions led to an increase in firing (Fig 1C), but AW to QW transitions led to a decrease in firing (Fig 1D). In some cases the morphology of the recorded neuron was revealed by applying the juxtacellular labeling method to the neuron recorded. The neuron shown in Fig 1B, C, D has been identified as a TC neuron based on its morphology (Fig 1E). The electrical activity of all morphologically identified LGN TC neurons (n=7) decreased their spontaneous firing during QW compared to AW (*p* < 0.001, Wilcoxon’s signed-rank test, Fig1F), but the strength of this modulation varied between neurons as shown on the distribution of their modulation indexes (Fig 1G).

**Figure 1.**
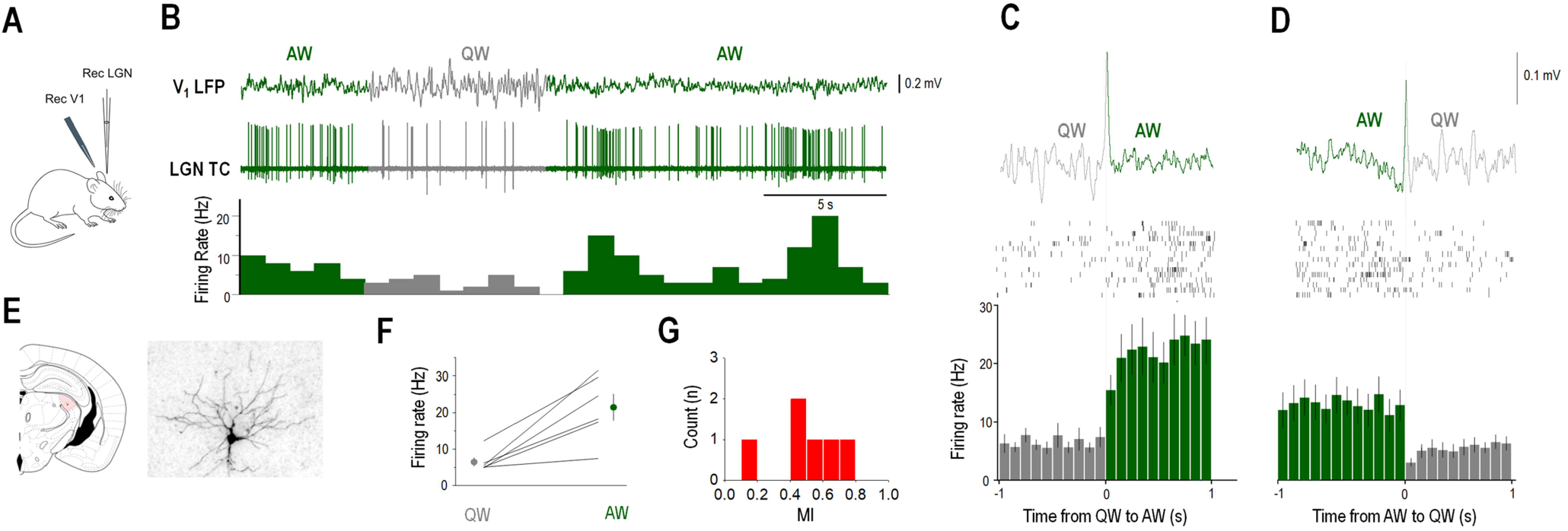
The baseline firing of identified LGN TC neurons is brain state dependent. (A) Schematics of the experimental setup. (B) Example of simultaneous V1 LFP and LGN unit recording from an awake head restrained mouse. Spontaneous state transitions are color coded for clarity (AW, active wakefulness, QW, quiet wakefulness). (B) Coronal brain section shows the morphology of the recorded neuron in the LGN. (C) QW to AW and (D) AW to QW transitions show prominent modulation of firing of this LGN TC neuron. State transition triggered LFP averages (top), rasters of the LGN TC neuron (middle) and PSTH (bottom), state transitions color coded as in (B). (F) Mean firing rate changes between QW and AW for all identified LGN TC neurons (n=7). (G) Distribution if modulation indexes (MI) for all identified LGN TC neurons (n=7).

To further explore the relationship between arousal and thalamic neuronal firing within the waking state we recorded the spontaneous firing of LGN neurons with simultaneous V_1_ and thalamic LFP and pupillometry. For the majority of these neurons, firing rates showed statistically significant positive correlation with pupil diameter (57 of 73 significant cells p < 0.05, random permutation test; Mean Pearson’s r = 0.411 ± 0.025, Fig 2AB). Fig 2C illustrates an example morphologically identified (Fig 2B) LGN TC neuron showing a positive correlation with pupil diameter. Note that the periods of dilated pupil coincide with AW states and high frequency tonic action potential output of the LGN TC neuron (Fig 2E), whereas the periods of constricted pupil with QW states and LTCP-mediated burst firing of the LGN TC neuron (Fig 2D). Thus periods of pupil constriction correspond to QW states as the V1 LFP shows a clear peak in the 5-9 Hz band (Fig 2D_2_), the cross-correlation of LFPs from V_1_ and LGN rhythmic activity at the two sites (Fig 2D_3_) and the autocorrelation of the TC neuron low frequency rhythmic burst firing (Fig 2D_4_). On the other hand, periods of pupil dilation correspond to AW states as the V1 LFP shows no peak in the 5-9 Hz band (Fig 2E_2_), the cross-correlation of LFPs from V_1_ and LGN is less rhythmic than during QW (compare with Fig 2D_3_) and the autocorrelation of the TC neuron high frequency tonic firing (Fig 2E_4_).

**Figure 2.**
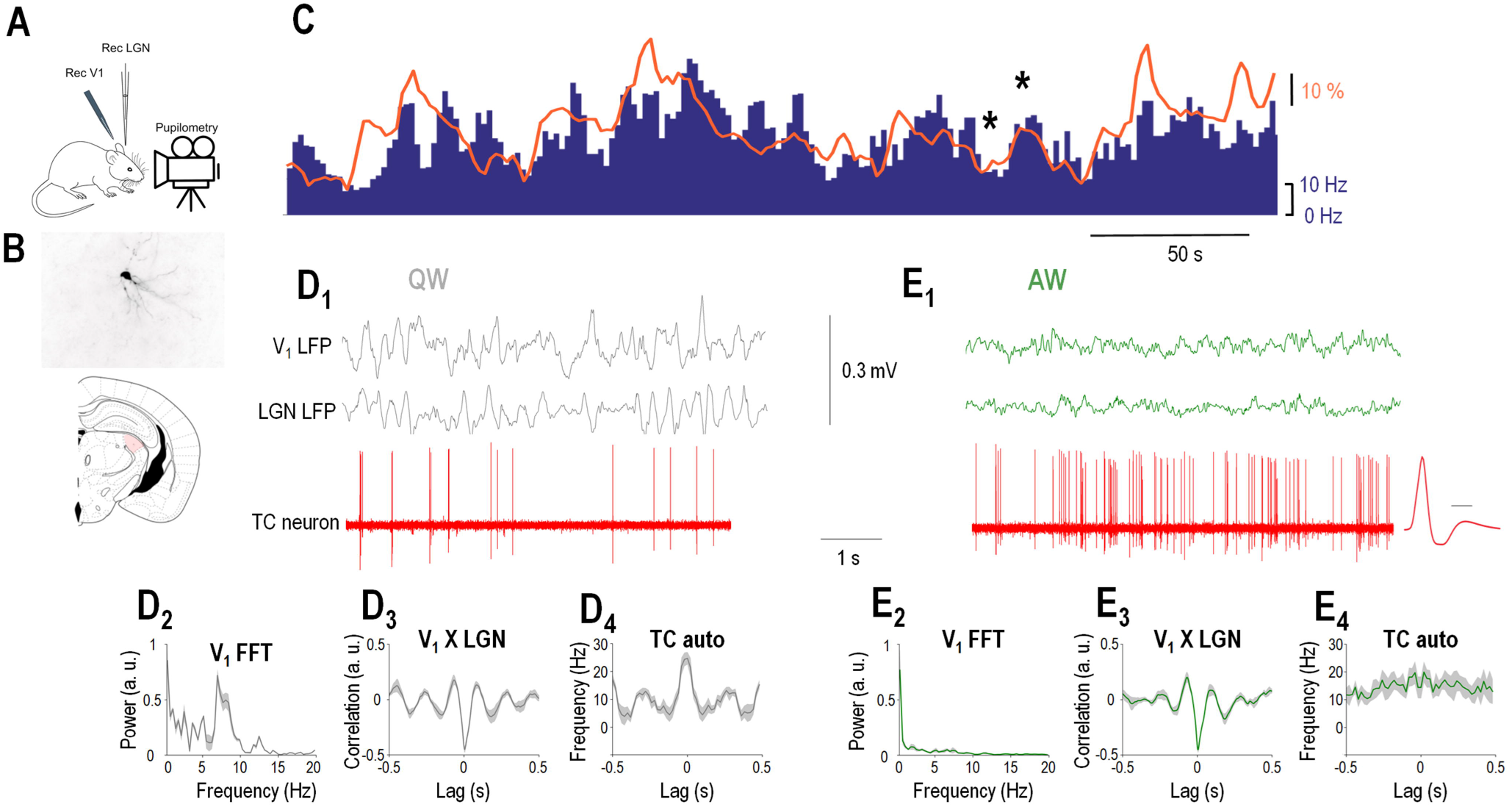
LGN TC neurons are positively correlated to arousal. (A) Schematics of the experimental setup. (B) Coronal brain section shows the morphology of the recorded neuron in the LGN. (C) Firing rate of an example juxtacellularly labeled LGN TC neuron (blue bars) increases when the simultaneously recorded pupil (orange line) is dilated, but decreases when the pupil is constricted. (D_1_) Simultaneously recorded visual cortical, thalamic LFP and single units of the TC neuron shown in (B and C) shown on a faster timebase during a period of constricted pupil (marked with an asterisk on C) corresponding to a state of QW. (D_2_) V_1_ power spectrum, (D_3_) cross correlation between V1 and LGN LFP and (D_4_) autocorrelation of the TC neuron during 10 consecutive periods of constricted pupil. (E_1_) Simultaneously recorded visual cortical, thalamic LFP and single units of the TC neuron shown in (B and C) plotted on a faster timebase during a period of constricted pupil (marked with an asterisk on C) corresponding to a state of AW. The average action potential waveform is shown in the right (time calibration: 0.5 ms). (E_2_) V_1_ power spectrum, (E_3_) cross correlation between V1 and LGN LFP and (E_4_) autocorrelation of the TC neuron during 10 consecutive periods of dilated pupil.

Strikingly, we found a smaller percentage of cells showing negative correlation to pupil diameter (16 of 73 state modulated thalamic neurons, 22%, p < 0.05, random permutation test; Mean Pearson’s r = −0.4 ±0.045). These neurons lacked classic LTCP mediated burtst firing, had narrower bi-phasic action potentials (Lorincz ML *et al*. 2009) and were thus classified as LGN interneurons. Fig 3B illustrates an example LGN interneuron showing a negative correlation with pupil diameter. Note that during the periods of constricted pupil characteristic of QW states (Fig 3C_2_ and C_3_) this LGN interneuron is more active than during AW (compare Fig 3C_4_ and 3D_4_). Also note that the activity of the LGN interneuron consists of tonic action potential output during both QW (Fig 3C_4_) and AW (Fig 3D_4_).

**Figure 3.**
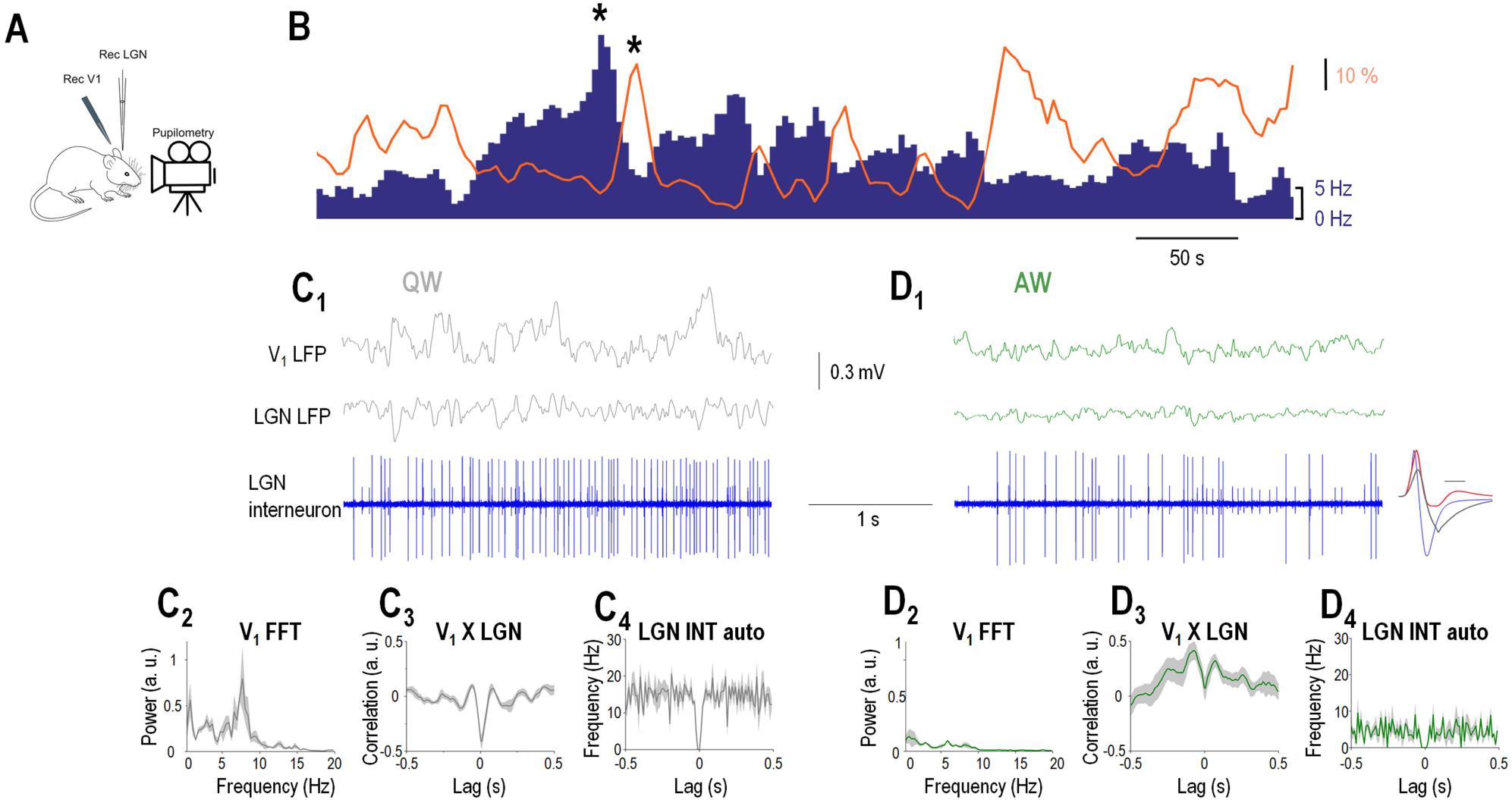
LGN interneurons are negatively correlated to arousal. (A) Schematics of the experimental setup. (B) Firing rate of an example LGN interneuron (blue bars) decreases when the simultaneously recorded pupil (orange line) is dilated, but increases when the pupil is constricted. (C_1_) Simultaneously recorded visual cortical, thalamic LFP and single units of the LGN interneuron shown in (B) plotted on a faster timebase during a period of constricted pupil (marked with an asterisk on C) corresponding to a state of QW. (C_2_) V_1_ power spectrum, (C_3_) cross correlation between V1 and LGN LFP and (C_4_) autocorrelation of the TC neuron during 10 consecutive periods of constricted pupil. (D_1_) Simultaneously recorded visual cortical, thalamic LFP and single units of the TC neuron shown in (B) plotted on a faster timebase during a period of constricted pupil (marked with an asterisk on B) corresponding to a state of AW. The average action potential waveform is shown in the right (blue, time calibration: 0.5 ms) with the average action potential of the TC neuron from Fig 2 overlaid for comparison. Note the presence of a second neuron (smaller spikes) not correlated with the pupil diameter, average action potential shown on the right in gray. (D_2_) V_1_ power spectrum, (D_3_) cross correlation between V1 and LGN LFP and (D_4_) autocorrelation of the TC neuron during 10 consecutive periods of dilated pupil.

**Figure 4.**
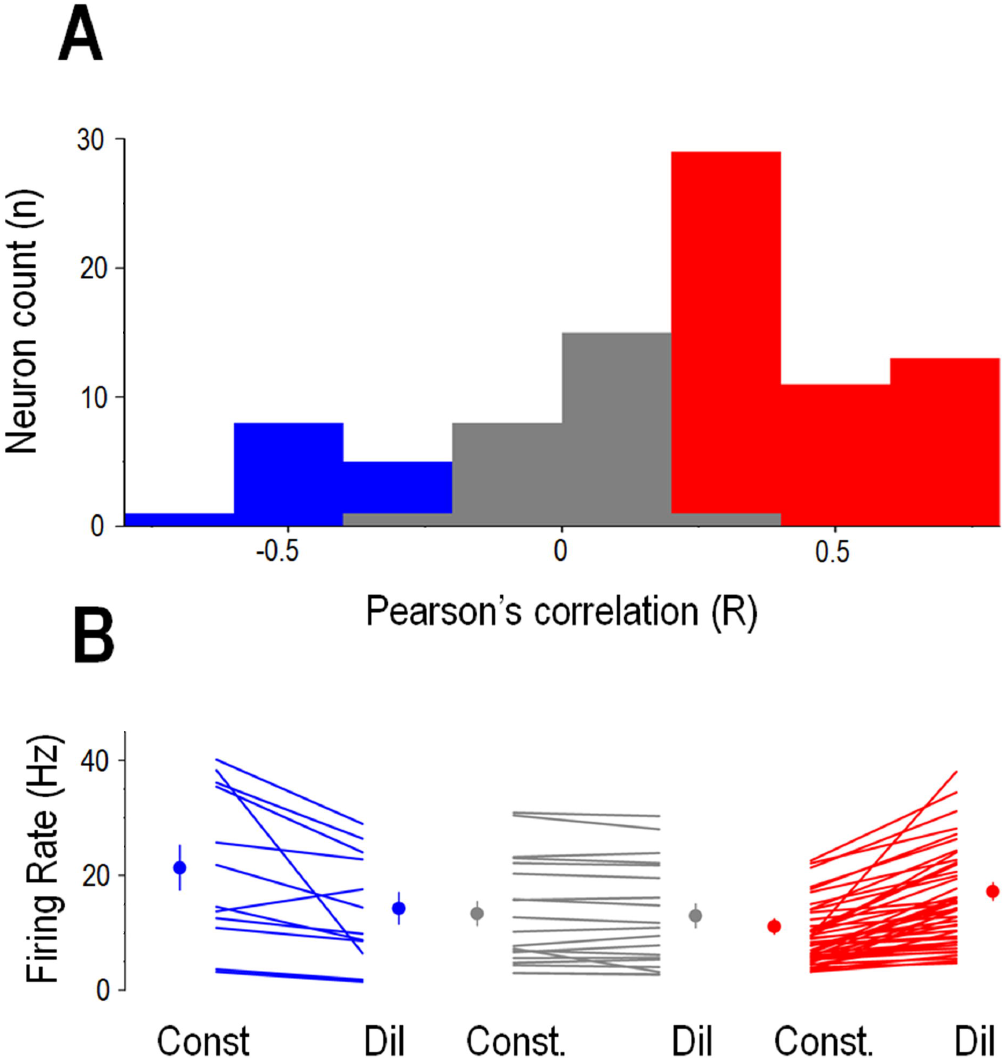
Arousal dependent activity in the LGN depends on neuronal identity. A) Mean firing rates of individual neurons during periods of constricted (Const) and dilated (Dil) pupil. LGN interneurons were negatively correlated (left, blue symbols), TC neurons were either uncorrelated (middle, gray symbols) or positively correlated (right, red symbols). Group averages are shown on the side of each group for the two states. (B) Distribution of pupil diameter/firing rate correlations, color codes same as in A.

All positively (n=57/98) correlated and non-correlated neurons (n=25/98) were identified as thalamocortical neurons based on morphological and/or physiological criteria, all negatively correlated neurons were physiologically identified as LGN interneurons (see Materials and Methods). Thus, LGN neurons in awake behaving mice form two groups where the spontaneous firing is subject to arousal dependent modulation being positively correlated with the pupil diameter in LGN TC neurons and negatively in LGN interneurons.

### The membrane potential of LGN TC neurons is correlated with brain states

To reveal the mechanisms underlying the state dependent fluctuation of thalamic neurons we performed intracellular recordings of LGN thalamocortical neurons (n=4) of awake mice while simultaneously monitoring the pupil diameter and recorded the LFP and multi-unit activity (MUA) in V_1_ (Fig 5A). In all the neurons recorded we found a clear-cut correlation between the membrane potential and pupil diameter (Fig 5B) such that periods of pupil constriction were associated with relatively hyperpolarized membrane potentials (~−65 mV) of thalamic neurons and burst firing in one of the neurons recorded (Fig 5A, bottom right). Importantly, these bursts were recorded at a membrane potential inconsistent with LTCP burst (Lorincz ML *et al*. 2009; Crunelli V et al. 2012; Crunelli V *et al*. 2018) and are therefore thought to represent high threshold bursts (Hughes SW *et al*. 2004; Lorincz ML et al. 2008; Lorincz ML *et al*. 2009; Crunelli V *et al*. 2012). Periods of pupil dilation, on the other hand, were associated with high frequency (Fig 5A and E) tonic action potential output and less hyperpolarized membrane potentials (~−55 mV, Fig 5A and F). When quantifying the correlation of the low pass filtered membrane potential and pupil diameter we found that the membrane potential of LGN TC neurons was lagging the changes in pupil diameter by ~5 ms (Fig 5C and D). Large amplitude (~5 mV) retinogeniculate EPSPs were not state dependent (Fig 5A inset, p>0.05, Wilcoxon’s signed-rank test). Thus, the state dependent action potential output of thalamic neurons can be accounted for by slow changes in neuronal membrane potential.

**Figure 5.**
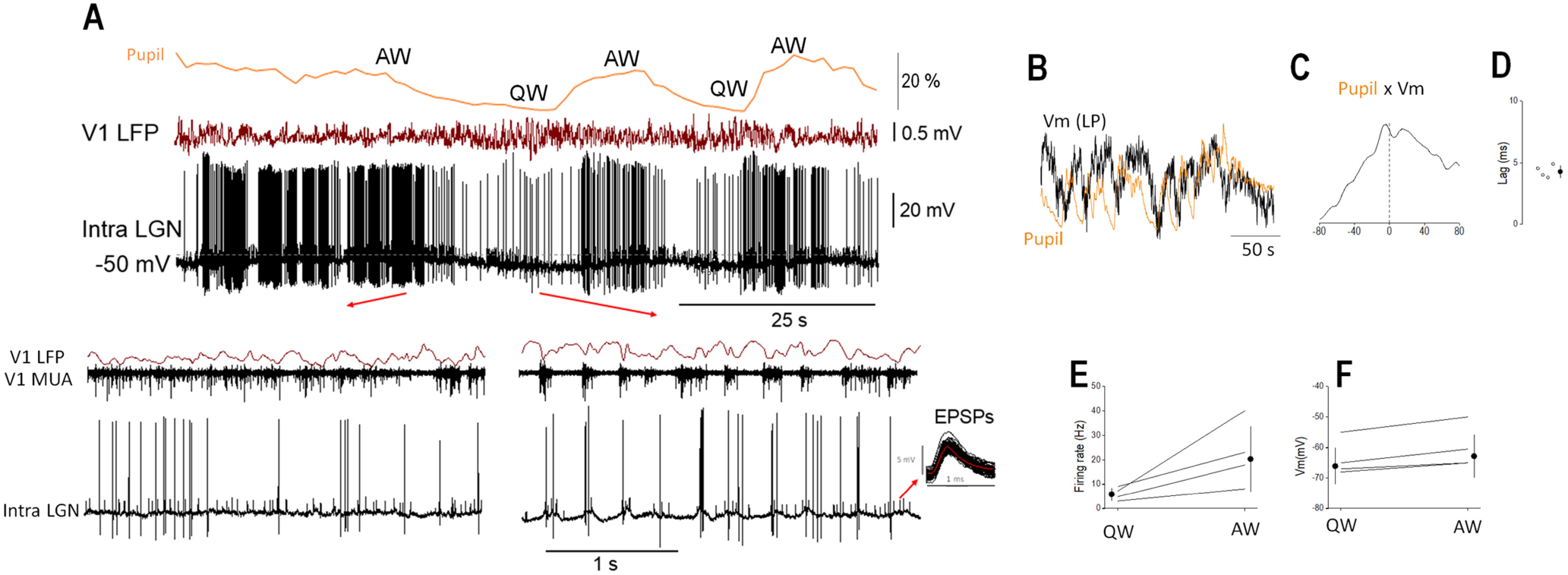
The membrane potential of LGN TC neurons is correlated with brain states. (A) Simultaneous pupilometry, cortical LFP, MUA and LGN Vm recording during state transitions in an awake head-restrained mouse. Quiet wake (QW) and active wake (AW) states are indicated above the pupilometry trace and shown on a faster timebase below. Note the large retinogeniculate EPSPs on the Vm recording. (B) Slow rhythmic fluctuations in the pupil diameter are correlated with the Vm of this LGN neuron as apparent on the low pass filtered Vm (Vm LP) overlaid on the pupilomerty trace. (C) Normailzed cross-correlation and quantification of Vm delay in respect to pupil diameter. Median firing rates (E) and Vm (F) for the two brain states.

### Inactivation of the visual cortex suppresses the brain state correlation of LGN neurons

As rhythmic synchronous LGN driven EEG activities are influenced by both cortical feedback and cholinergic inputs to the thalamus (Lorincz ML *et al*. 2009) we monitored the effects of inactivating the cortical feedback from V_1_ to the LGN and monitor its effect on the arousal dependent activity of LGN neurons. The firing rate of individual neurons before and after V1 inactivation varied considerably (Fig 6C) but did not reach statistical significance as a group (*p* □ 0.05, Wilcoxon’s signed-rank test, n=7, Fig 6D). When comparing the correlation of thalamic single units and pupil diameter before and after V1 inactivation (Fig 6B and C) we found a significant decrease (*p* < 0.001, Wilcoxon’s signed-rank test, n=7) suggesting that corticothalamic input from V1 is at least partly responsible for the state dependent activity in LGN TC neurons.

**Figure 6.**
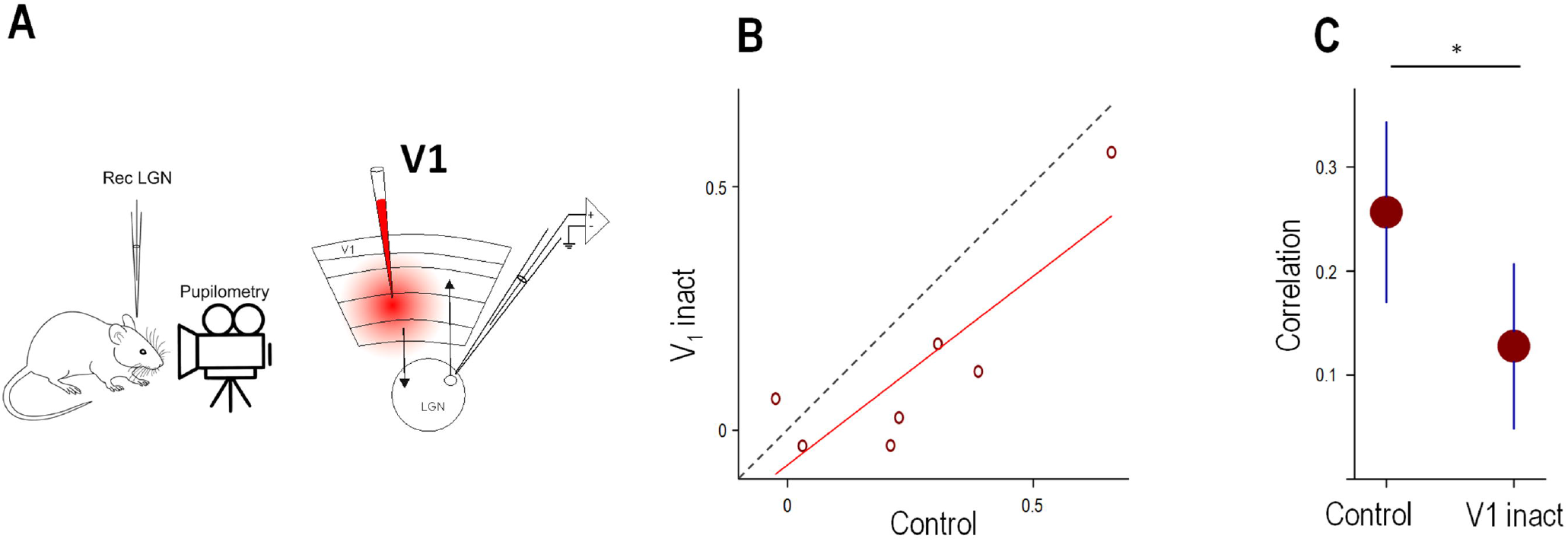
V1 inactivation decreases the correlation of LGN neuron firing and pupil diameter. (A) Schematics of the experimental setup. (B) Scatter plot comparing the firing rate/pupil diameter correlation before (Control) and after the pharmacological inactivation (V1 inact) of the ipsilateral V1. (C) Mean control and inactivation correlations. (D) Scatter plot comparing the firing rates of LGN TC neurons before (Control) and after the pharmacological inactivation (V1 inact) of the ipsilateral V1. (E) Mean control and inactivation firing rates.

## DISCUSSION

Thalamocortical dynamics are essential for sensory coding, cognition and brain rhythms. Whereas cortical function in general and visual cortical activity in particular are well known to be modulated by the level of arousal in awake animals (McGinley MJ, M Vinck, *et al*. 2015) the arousal dependent activity of reciprocally connected visual thalamus is more controversial. Here, in awake behaving mice we found that i) the baseline LGN neuronal activity correlates with arousal during the awake state, ii) the polarity of this correlation is cell type specific, with LGN TC neurons being positively, and LGN interneurons negatively correlated with arousal and iii) this state dependent activity at least partly originates from cortical feedback.

LGN activity has long been known to differ during wakefulness compared to sleep (Hirsch JC et al. 1983), during alert and inattentive states (Bezdudnaya T *et al*. 2006), with a subset of bursting LGN TC neurons playing an important role in generating, while LGN interneurons in providing tonic firing TC neurons with temporally precise phasic inhibition during individual cycles of the alpha rhythm (Lorincz ML *et al*. 2009). Some results suggested that LGN spontaneous and visually evoked firing rates are not different between immobile and running mice, whereas V1 activity was markedly different between the two states (Niell CM and MP Stryker 2010). This controversy might arise from differences in the definition of brain states: whereas locomotion is only associated with alert wakefulness, both quiet and alert wakefulness can occur during immobility (Vinck M *et al*. 2015). Pupil diameter is an excellent proxy for a general neuromodulatory tone and brain states (Reimer J et al. 2016) and its combination with V1 LFP level of synchronization as used in the present study provides a more accurate state definition.

Our results reveal an arousal dependent spontaneous activity in the majority of LGN neurons of awake behaving animals. A recent study found that TC neurons of the primary and higher order somatosensory thalamus also show state dependent activity when comparing activities recorded during quiet wakefulness and sleep (Urbain N et al. 2019). Notably, the polarity of the correlation between neuronal activity and arousal, quantified by monitoring the pupil diameter depends on neuronal identity within the LGN. Specifically, whereas TC neurons, the dominant cell type in the LGN tend to increase their activity during active wakefulness (desynchronized V1 LFP and dilated pupil) LGN interneurons, that are key for intimately controlling the timing of TC neuron firing in awake animals (Lorincz ML *et al*. 2009), behave in an opposite manner showing decreased activity during AW and increased activity during QW (synchronized V1 LFP and constricted pupil). Taken together, the increase in firing in most TC neurons during AW is accompanied by a decrease in inhibition derived from local interneurons (Acuna-Goycolea C et al. 2008; Lorincz ML *et al*. 2009). This may have implications in both the spontaneous activity of LGN TC neurons and their sensory coding. The decrease in firing in LGN interneurons during AW does not necessarily mean that the net inhibition in TC neurons is decreased as some TRN neurons, the sources of a much larger inhibitory conductance (Bal T et al. 1995) are known to fire at much higher rates during AW (Gardner RJ et al. 2013; Halassa MM et al. 2014). The significance of this arousal dependent modulation of inhibition derived from local interneurons in rhythmic brain activity and sensory coding needs to be addressed by future studies.

What mechanisms are responsible for the differential arousal-dependent activity of LGN TC and interneurons? Our intracellular recordings in TC neurons reveal that during AW states the membrane potential of LGN TC neurons is less hyperpolarized than during QW, providing an explanation for the increases in firing rate during AW in LGN TC neurons. The firing mode of these neurons also changed from tonic to burst firing in the QW state, and in one case (Fig 5) these bursts were characterized by a membrane potential polarization and interspike-interval inconsistent with LTCP mediated burst firing, suggesting they are high-threshold bursts similar to the ones recorded in the feline LGN (Hughes SW *et al*. 2004; Lorincz ML *et al*. 2009) or somatosensory thalamus of lightly anesthetized mice (Crunelli V *et al*. 2012). Some of our extracellular recordings during QW clearly show LTCP like bursts (FIG 2D_1_), suggesting that high-threshold bursting is a property of a subset of LGN TC neurons as it was described in the cat LGN (Hughes SW *et al*. 2004; Lorincz ML *et al*. 2008; Lorincz ML *et al*. 2009). Importantly, we revealed a tight correlation between TC neuron membrane potential and pupil diameter (Fig 5B and C) suggesting that the arousal dependent membrane potential might originate from differential neuromodulation during AW and QW states. Multiple brain sources could contribute to this modulation, the most evident ones being brainstem cholinergic (de Lima AD et al. 1985) and corticothalamic fibers (Wilson JR et al. 1984). Indeed, upon V1 inactivation we found a prominent suppression of the arousal dependency of LGN TC neurons suggesting that corticothalamic feedback is at least partly responsible for this phenomenon. If ACh is indeed involved in this modulation it could explain the differential arousal related behavior revealed here in TC neurons and interneurons, respectively, because whereas TC neurons are depolarized (McCormick DA 1992; Lorincz ML *et al*. 2008) LGN interneurons are hyperpolarized by ACh (McCormick DA and HC Pape 1988). Thus, an increase in cholinergic tone during AW could explain both the increased TC neuron and decreased LGN interneuron firing. Neurons in the brainstem cholinergic nuclei show pronounced arousal dependent activity (Steriade M et al. 1990) making them a good candidate for mediating these effects. Taken together our results show that LGN membrane potential and action potential output are dynamically linked to arousal dependent brain states in awake mice and this fact might have important functional implications.

